# Trends of genetic changes uncovered by Env- and Eigen-GWAS in wheat and barley

**DOI:** 10.1101/2020.11.27.400333

**Authors:** Rajiv Sharma, James Cockram, Keith A. Gardner, Joanne Russell, Luke Ramsay, William TB Thomas, Donal M. O’Sullivan, Wayne Powell, Ian J. Mackay

## Abstract

The process of crop breeding over the last century has delivered new varieties with increased genetic gains, resulting in higher crop performance and yield. However in many cases, the underlying alleles and genomic regions that have underpinned this success remain unknown. This is due, in part, to the difficulty in generating sufficient phenotypic data on large numbers of historical varieties to allow such analyses to be undertaken. Here we demonstrate the ability to circumvent such bottlenecks by identifying genomic regions selected over 100 years of crop breeding using the age of a variety as a surrogate for yield. Using ‘environmental genome-wide association scans’ (EnvGWAS) on variety age in two of the world’s most important crops, wheat and barley, we found strong signals of selection across the genomes of our target crops. EnvGWAS identified 16 genomic regions in barley and 10 in wheat with contrasting patterns between spring and winter types of the two crops. To further examine changes in genome structure in wheat and barley over the past century, we used the same genotypic data to derive eigenvectors for deployment in EigenGWAS. This resulted in the detection of seven major chromosomal introgressions that contributed to adaptation in wheat. The deployment of both EigenGWAS and EnvGWAS based on variety age avoids costly phenotyping and will facilitate the identification of genomic tracts that have been under selection during plant breeding in underutilized historical cultivar collections. Our results not only demonstrate the potential of using historical cultivar collections coupled with genomic data to identify chromosomal regions that have been under selection but to also guide future plant breeding strategies to maximise the rate of genetic gain and adaptation in crop improvement programs.

**Significance Statement:** 100 years of plant breeding have greatly improved crop adaptation, resilience, and productivity. Generating the trait data required for these studies is prohibitively expensive and can be impossible on large historical traits. This study reports using variety age and eigenvectors of the genomic relationship matrix as surrogate traits in GWAS to locate the genomic regions that have undergone selection during varietal development in wheat and barley. In several cases these were confirmed as associated with yield and other selected traits. The success and the simplicity of the approach means it can easily be extended to other crops with a recent recorded history of plant breeding and available genomic resources.

## Introduction

In the last century, significant improvements in yield and quality have been reported in almost all crop species as a result of plant breeding driven by market demand (1). However, the growing demand for food, feed and fibre to meet the expanding global human population requires an acceleration in the pace of crop genetic improvement (2). Identification of the genetic loci responsible for these changes will help accelerate the genetic gains required to meet future food security needs, via their incorporation in marker assisted selection breeding strategies (3). Over the last decade, genome wide association studies (GWAS) has become a prominent method for genetic analysis in plants (4). In crops, GWAS require trait data on large collections of varieties or accessions, which is typically expensive to collect and can therefore result in underpowered studies with relatively low numbers of lines (5, 6). An alternative is to exploit the availability of historical data, such as that collected during varietal development programmes.

For almost every major crop, yield is the most important breeding target. Breeding programmes invest large amounts of resources into realising the incremental genetic gains in yield that are required for continual varietal improvement. Accordingly, the process of developing new crop varieties involves rigorous screening in large multi-location and multi-environmental trials over several years. Large historical phenotypic data sets from such trials have been successfully employed for GWAS in the past (7), and in several cases have identified the functional genes underlying the genetic control of the investigated traits (8–11). However, the availability of seed for variety collections with appropriate trait data is not common for many crops. Alternatively, seed of historical varieties may be available, but the associated trait data may be lost or disjointed. In both cases, the cost of collecting *de novo* trait data can be prohibitive. In many cases however, the release date, subsequently termed here ‘age’, of varieties is known. Given that in most crops, the breeding process has improved the genetic potential of key agronomic traits over time, variety age can be used as a surrogate measure of merit and mapped in GWAS. This approach, in which environmental or non-genetic variables are treated as traits in GWAS to map loci associated with those variables, has been termed EnvGWAS (12). For many crops, the predominant genetic change over time has been to increase yield (e.g.(13)), and the age of a variety may function directly as a surrogate for yield, although loci detected may also be associated with other temporal changes. EnvGWAS on variety age can also be regarded as a simple genome-wide test for genetic loci under directional selection, which may be subsequently associated with traits. This approach may also provide a way of identifying alleles associated with adaptation (14) which otherwise have been difficult to detect. Finally, EnvGWAS can be a cost effective strategy since it can access large pre-existing datasets but is not dependent on historical or *de novo* trait data.

A related approach requiring no trait data is EigenGWAS (15). Using genotypic data alone, the singular value decomposition of the genomic relationship matrix provides loadings (eigenvectors) for each variety on each eigenvalue of the matrix. For the largest eigenvalues, these loadings are then treated as independent traits for GWAS. Significant associations with any particular component highlight genomic regions or markers of greatest importance for that eigenvalue, and therefore the potential major drivers of population structure. Subsequent study of varieties differing in these regions may also be interpretable in terms of drivers of adaptation. EigenGWAS and EnvGWAS have recently been used to study diversity among maize landraces and identify lines and traits suitable for downstream analysis without large scale phenotyping (12).

In this study, we demonstrate for the first time the utility of treating variety age as a surrogate trait for crop productivity when combined with EnvGWAS and EigenGWAS to identify target regions and quantitative trait loci (QTL) underpinning genetic improvements in crop performance that have occurred during modern plant breeding. This is a powerful but cost effective method that does not require extensive trait data or complex software. We demonstrate the utility of these complementary approaches by: i) using EnvGWAS on variety age to identify loci responsible for genetic improvement in four complimentary datasets of modern winter and spring types of wheat (*Triticum aestivum*) and barley (*Hordeum vulgare*) from the United Kingdom (UK) and Brazil. ii) Validating the results from (i) by GWAS on subsets of these varieties for which historic yield data were also available. iii) Evaluating the temporal changes of allelic state at the loci identified. iv) Performing EigenGWAS on the same four datasets. EigenGWAS compliments EnvGWAS in that it too does not required trait data and may also identify genomic regions that have undergone selection. However, unlike EnvGWAS, it does not explicitly search for regions associated with variety age and is more likely to detect features associated with local adaptation, which may change little in frequency over time. As far as we are aware, no EnvGWAS analysis has been published in plants for which variety age has been used as a trait. The combination of EnvGWAS with EigenGWAS used here provides insights into the recent breeding history and population structure of two of the world’s most important crops, and highlights the effectiveness and simplicity of these approaches to study recent selection history without the requirement for phenotype data.

## Results

### Year of variety release as a surrogate measure for yield

The Pearson correlations between historical yield data and age of variety were calculated for the subsets of 192 UK wheat and 197 UK barley varieties for which historical yield data were available (***SI Appendix*, Fig. S1**). High correlations between yield and year of release (range 0.896 – 0.974) were found in both UK data sets. This confirms year of release could be used as a good measure of genetic progress in UK wheat and barley yield potential. No historical yield data for the Brazilian wheat panel were available.

### EnvGWAS for variety age

*EnvGWAS wheat*. Using variety age for EnvGWAS in the UK winter wheat panel (*n=*404) identified thirteen significant (-log_10_ (*p*) >4.0) genomic regions, of which four loci were found to be highly significant (-log_10_ (*p*) >6.0), located on chromosomes 1A, 2A, 2D and 6A (**Fig. 1A, Table 1, *SI Appendix*, Table S1**). For Brazilian spring wheat (*n=*355), three significant genetic loci were detected, two on chromosome 2B (251 cM, 318 cM) and one on 5A (710 cM), none of which were identified in the UK winter wheat panel (**Fig. 1B, Table 1, *SI Appendix*, Table S1)**.

*EnvGWAS Barley*. We identified three highly significant genetic loci in the winter barley panel (*n=*297), and seven in the spring barley panel (*n=*406) (**Table 1; Fig. 1C-D**); a summary of the associated markers is listed in ***SI Appendix*, Table S2**. Two significant loci were identified in both barley panels (chromosome 3H, ∼68-70 cM; 5H, ∼20 cM) (**Fig. 1** and **Table S2**). Subsequently, EnvGWAS was performed on the combined winter and spring panels (*n*=704), identifying the same four significant loci we identified in the spring panel alone (***SI Appendix*, Fig. S2A, Table S2** and **Table 1**). We repeated the analysis using seasonal growth habit (‘spring’ or ‘winter’ types) as a covariate, without any major changes in results (***SI Appendix*, Fig. S2B**). In addition, we performed GWAS on seasonal growth habit itself, identifying three major genetic loci on the long arms of chromosomes 1H, 4H and 5H (***SI Appendix*, Fig. S2C**), corresponding to major flowering time and vernalization genes known to be the major determinants of winter and spring seasonal growth type (*PPD-H2* on chromosome 1H, *VRN-H2* on 4H and *VRN-H1* on 5H) (16, 17). EnvGWAS for variety age was then repeated with these QTL as covariates (***SI Appendix*, Fig. S2D**). The most significant results mainly on chromosome 5H from the analyses with and without covariates changed little. However, the magnitude of other significant peaks differed, such as the locus on chromosome 1H.

**Table 1.** Summary of the singificant hits detected by EnvGWAS on variety age. Details in SI Appendix, Table S1 & Table S2.

| Pop-name | SNP-name | Chrom | Position (cM) | Ref-allele | Ref-Allele-F | -log(p) | Effects |
| --- | --- | --- | --- | --- | --- | --- | --- |
| Winter-Wheat | wsnp_Ex_c572_1A | 1A | 221.0 | A | 0.50 | 6.01 | -7.61 |
|  | Kukri_c18109_1B | 1B | 350.0 | A | 0.92 | 4.62 | 11.64 |
|  | Excalibur_c153_2A | 2A | 20.0 | A | 0.66 | 6.31 | 9.50 |
|  | RFL_Contig403_2A | 2A | 62.0 | A | 0.65 | 5.24 | 8.32 |
|  | BS00071630_5_2A | 2A | 87.0 | A | 0.66 | 6.18 | 9.26 |
|  | IACX6178_2A | 2A | 158.0 | A | 0.66 | 6.18 | 9.26 |
|  | BS00022799_5_2D | 2D | 33.0 | A | 0.66 | 6.31 | 9.50 |
|  | BobWhite_rep_5B | 5B | 381.0 | A | 0.13 | 4.31 | 6.94 |
|  | BS00021901_5_5D | 5D | 180.0 | T | 0.85 | 5.04 | 9.58 |
|  | BS00022120_5_6A | 6A | 190.0 | T | 0.83 | 8.11 | 12.87 |
|  | Kukri_c16404_6B | 6B | 322.0 | A | 0.06 | 4.06 | 10.33 |
|  | Kukri_c67076_7A | 7A | 383.0 | A | 0.14 | 4.29 | 8.48 |
|  | BobWhite_c429_7B | 7B | 236.0 | A | 0.94 | 4.88 | -12.92 |
| Spring-Wheat | Ku_c5725_892_2B | 2B | 251.0 | A | 0.49 | 4.44 | -7.35 |
|  | RFL_Contig484_2B | 2B | 318.0 | T | 0.76 | 4.20 | -9.34 |
|  | RAC875_c8642_5A | 5A | 710.0 | A | 0.08 | 4.51 | -13.21 |
| Winter-Barley | JHI-Hv50k-2011_3H | 3H | 68.7 | A | 0.29 | 4.74 | -1.95 |
|  | JHI-Hv50k-2011_3H | 3H | 124.5 | C | 0.64 | 4.22 | 1.71 |
|  | JHI-Hv50k-2011_5H | 5H | 19.2 | A | 0.73 | 5.92 | -1.87 |
| Spring-Barley | JHI-Hv50k-2011_1H | 1H | 51.0 | A | 0.41 | 4.08 | -3.07 |
|  | SCRI_RS_1486_2H | 2H | 0.0 | A | 0.42 | 5.17 | -2.59 |
|  | JHI-Hv50k-2011_3H | 3H | 1.7 | C | 0.22 | 4.40 | 3.69 |
|  | JHI-Hv50k-2011_3H | 3H | 77.7 | C | 0.95 | 4.52 | -4.42 |
|  | JHI-Hv50k-2011_5H | 5H | 20.5 | C | 0.12 | 4.90 | 3.38 |
|  | 12_30230_6H | 6H | 53.1 | A | 0.88 | 5.22 | 4.45 |
|  | JHI-Hv50k-2011_7H | 7H | 7.8 | A | 0.93 | 5.40 | 5.37 |
| Spring&Winter- | JHI-Hv50k-2011_2H | 2H | 0.0 | C | 0.74 | 4.17 | -2.15 |
|  | JHI-Hv50k-2011_2H | 2H | 20.3 | C | 0.92 | 5.74 | -2.86 |
|  | JHI-Hv50k-2011_3H | 3H | 45.2 | C | 0.92 | 4.15 | 3.07 |
|  | JHI-Hv50k-2011_3H | 3H | 68.7 | C | 0.14 | 7.13 | -4.41 |
|  | JHI-Hv50k-2011_3H | 3H | 126.6 | C | 0.80 | 4.29 | 3.64 |
|  | JHI-Hv50k-2011_5H | 5H | 19.2 | C | 0.82 | 7.67 | -3.43 |
|  | JHI-Hv50k-2011_5H | 5H | 105.0 | A | 0.09 | 4.51 | 3.53 |
|  | 11_20546_5H | 5H | 160.7 | A | 0.89 | 4.70 | -2.94 |
|  | JHI-Hv50k-2011_7H | 7H | 3.8 | C | 0.05 | 5.56 | -4.65 |

**Figure 1.**
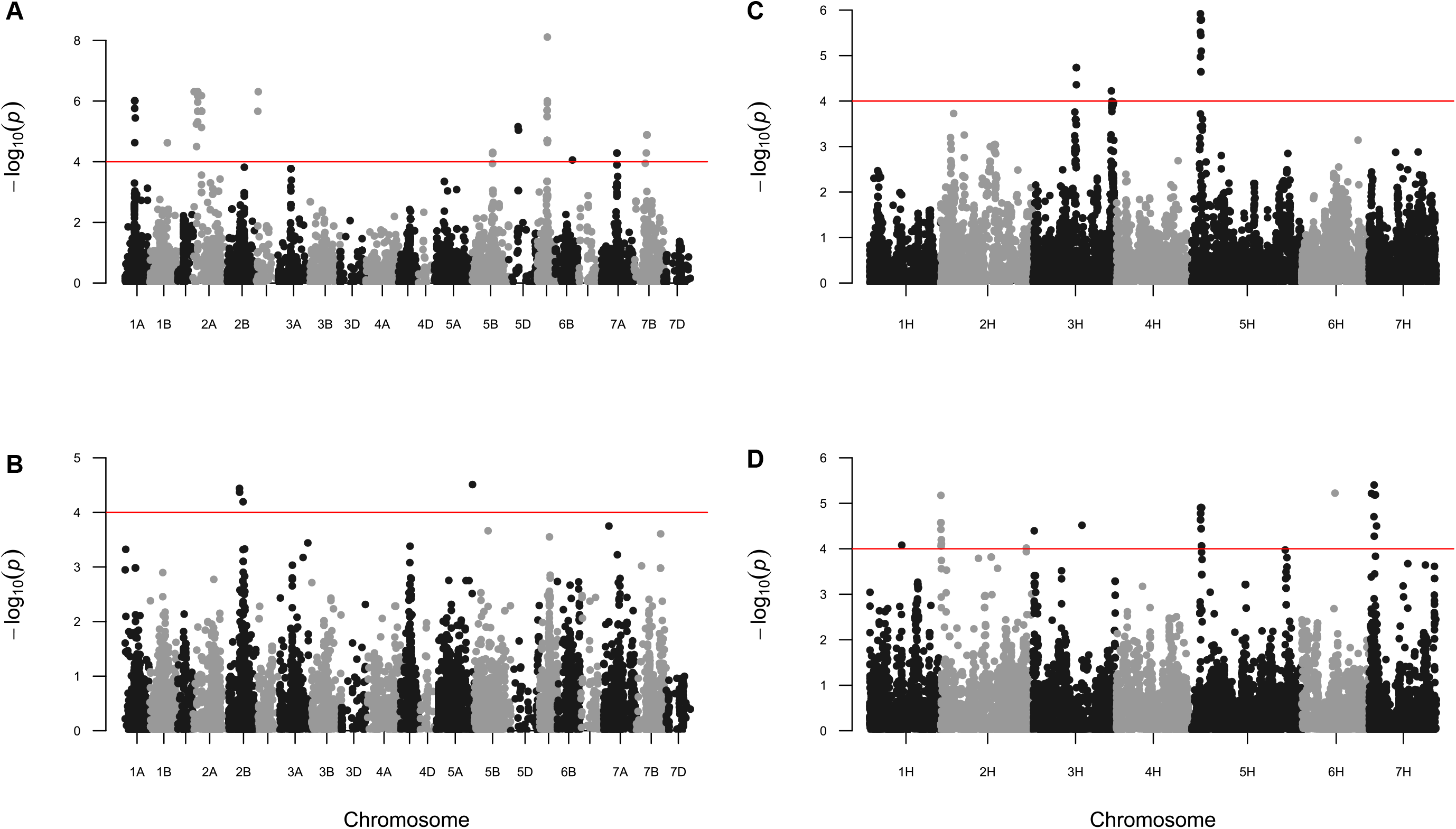
Wheat EnvGWAS for variety age. Manhattan plots of the four panels are shown. On the x-axis genetic positions are based on the consensus map (41) are displayed for (*A*) UK winter wheat and (*B*) Brazilian spring wheat panels; a pseudo-genetic map positions that relates to the physical positions (40) of the UK winter (*C*) and spring (*D*) barley panels are shown. On the y-axis − log_10_(*p*)-values are displayed. The red line indicates the threshold value of the significance corresponding to − log_10_(*p*) = 4.

### Validation of EnvGWAS based on trait analysis and a multi-founder experimental population

To validate the EnvGWAS analyses, we performed GWAS on the subset of 192 UK winter wheat varieties for which historical yield data were available together with EnvGWAS on variety age for direct comparison of the results. In this subset, we found that GWAS for yield identified the same genomic region on chromosome 1A (**Fig. S3A**) as was detected by EnvGWAS for variety age (***SI Appendix*, Fig. S3, Table S1**). This is the same region that we identified in EnvGWAS for variety age in the complete set of 404 UK wheat varieties. Interestingly, while the chromosome 5A QTL was detected with low-significance (-log_10_ (*p*) = 4.45) by GWAS on yield, it was not identified using EnvGWAS on variety age. In addition, EnvGWAS analysis of variety life-span also detected a locus on chromosome 1B that was not detected in any other of our analyses.

Similarly, EnvGWAS on variety age and GWAS on yield was repeated using the subset of 197 winter and spring barley varieties for which historic yield data was available, detecting highly significant hits (-log_10_ (*p*) > 4.0) on chromosome 5H for variety age, variety life-span and yield, using seasonal growth habit as a covariate (***SI Appendix*, Fig. S4, Table S2**). Although not identified in the larger panel of 703 varieties, analysis of our subset of 197 lines consistently identified a highly significant genetic locus on the short arm of chromosome 3H for variety age, variety life-span and yield. An additional peak was detected with EnvGWAS for variety life-span on the long arm of chromosome 2H.

To further validate our EnvGWAS findings, we analysed data from a 16 founder wheat multiparent advanced generation inter cross (MAGIC) population consisting of 550 recombinant inbred lines generated by inter-crossing 16 wheat varieties released between 1935 to 2004 (18). We found that the four major genomic regions previously identified by EnvGWAS of variety age on chromosomes 1A, 2A, 2D and 6A were also significant in the MAGIC population for several yield and grain related traits, height, and yellow-rust resistance (**Table 2**). Further details of the 213 agronomic and disease resistance traits analysed and the corresponding significance levels are listed in (***SI Appendix*, Table S3**).

**Table 2:** Collocation of significant loci (-log(p)>3) in MAGIC with the three major winter wheat GWAS peaks. Collocation used 35K and 90K physical maps.

| **Trait (MAGIC coding)** | **Trait decription** | **Region** |
| --- | --- | --- |
| GLA_on_31_03_17_Year1 | Grain Length and Area | 1A |
| waxiness_leaf_score_Year1 | Waxiness | 1A |
| FL_length_x_width_Year1 | Flag-leaf length | 1A |
| Pigmentation_score_Year1 | Pigmentation-score | 1A |
| Height_FL_to_ear_base_Year1 | Height | 1A |
| Spikelets_paired_frequency_in_20_Year1 | Spikelets-pair-freq | 1A |
| Spring_type_Year1 | Spring-type | 1A |
| Yellow_rust_on_11_05_17_Year1 | Yellow-rust | 2A |
| GLA_on_20_04_17_Year1 | Grain Length and Area | 2A |
| Chlorosis_score_Year1 | Chlorosis-score | 2A |
| GS65_DAS_Year1 | Heading | 2A |
| FL_width_Year1 | Flag-leaf width | 2A |
| Yield_Year1 | Yield | 2A |
| GLA_on_07_03_17_Year1 | Grain Length and Area | 6A |
| waxiness_leaf_score_Year1 | Waxiness | 6A |
| FL_angle_Year1 | Flag-leaf angle | 6A |
| FL_length_Year1 | Flag-leaf length | 6A |
| FL_length_x_width_Year1 | Flag leaf area | 6A |
| FL_length_width_ratio_Year1 | Flag leaf length/width ratio | 6A |
| Height_to_FL_Year1 | Height | 6A |
| Height_to_ear_base_Year1 | Height | 6A |
| Height_to_ear_tip_Year1 | Height | 6A |
| Ear_length_Year1 | Ear-length | 6A |
| Height_FL_to_ear_base_Year1 | Height | 6A |

### Allele-shift over time

To illustrate the changes in allele frequency present in our variety collections over time, we generated rocket plots (***SI Appendix*, Fig. S5–S8**) for the major genomic regions identified by EnvGWAS on variety age (***SI Appendix*, Table S4-S7**). Different patterns and intensity of selection were evident across chromosomal regions over time. For wheat, these fell into three broad classes: (1) Late introduction of ‘modern’ alleles followed by a rapid increase in frequency (***SI Appendix*, Fig. S5A**), (2) retention of both ‘modern’ and ‘old’ alleles at similar frequency across time (e.g. ***SI Appendix* S5E**), (3) relatively early introduction of the ‘modern’ allele, followed by its retention at low frequency (e.g. ***SI Appendix* S5F**). Details of the alleles-shifts examples are provided in the Supplementary Notes. In barley, the rocket plots illustrated both gradual and rapid shifts in allele frequency at the genomic regions identified by EnvGWAS on variety age (***SI Appendix*, Fig. S5I-N**). For example, for the UK spring barley genetic locus on chromosome 7H (∼8.8 Mbp), only one allele was present until 1992 (***SI Appendix*, Fig. S5N and Table S6**), after which the ‘modern’ allele remained at low frequency, even among modern varieties. A genomic region on chromosome 5H which was identified separately in winter and spring barley displays a pattern where the ‘modern’ allele is introduced in 1986, after which both alleles are found at intermediate frequencies among the most recent varieties in winter barleys. However, modern spring barleys were predominantly of ‘modern’ allele type.

### EigenGWAS scans

While EnvGWAS allowed us to use variety age to investigate the genomic regions underlying QTL for yield and adaptation, we hypothesised that the complementary method, EigenGWAS, would allow us to detect changes in larger scale structural variants in our target crop genomes over time.

After determining the first ten PCs in each of our UK and Brazilian wheat populations **(*SI Appendix*, Table S8)**, EigenGWAS detected numerous significant hits (N=11567 SNPs with -log_10_ (*p*) >4.0) (**Fig. 2** & ***SI Appendix*, Table S9**). Seven genetic loci distributed on chromosomes 1A, 1B, 2B, 5B, 6A and 6B were found to be significant with multiple PCs, as well as within the Brazilian (spring) and UK (winter) panels **(Fig. 2)**. These loci corresponded to major chromosomal introgressions from related cereal species into wheat (***SI Appendix*, Table S9**). For instance, the 1B locus co-locates with the chromosome 1B/1R introgression from rye (*Secale cereale*), which is known to regulate multiple traits including disease resistance and yield (19, 20). We identified an additional seventeen putative introgressions that were supported by a recent introgression survey by (21), along with another 58 novel putative introgressions (***SI Appendix*, Table S9**). Among these novel putative introgressions were regions on chromosome 5A, depicted in **Fig. 2** as 5A_2 and 5A_3, which displayed amongst the most significant hits across the UK and Brazilian wheat data sets and multiple PCs. Interestingly, two highly significant genomic regions (1A_2 and 5A_5) identified by EigenGWAS on PC2 in winter wheat were also detected by GWAS on yield in the validation data set (***SI Appendix*, Table S13**). In addition, three genomic regions (5B_2, 6A_1 and 7B_1) identified in the winter wheat EigenGWAS analysis were also detected in EnvGWAS on variety age, suggesting both approaches are not exclusively identifying different genomic regions (***SI Appendix*, Table S13**).

**Figure 2.**
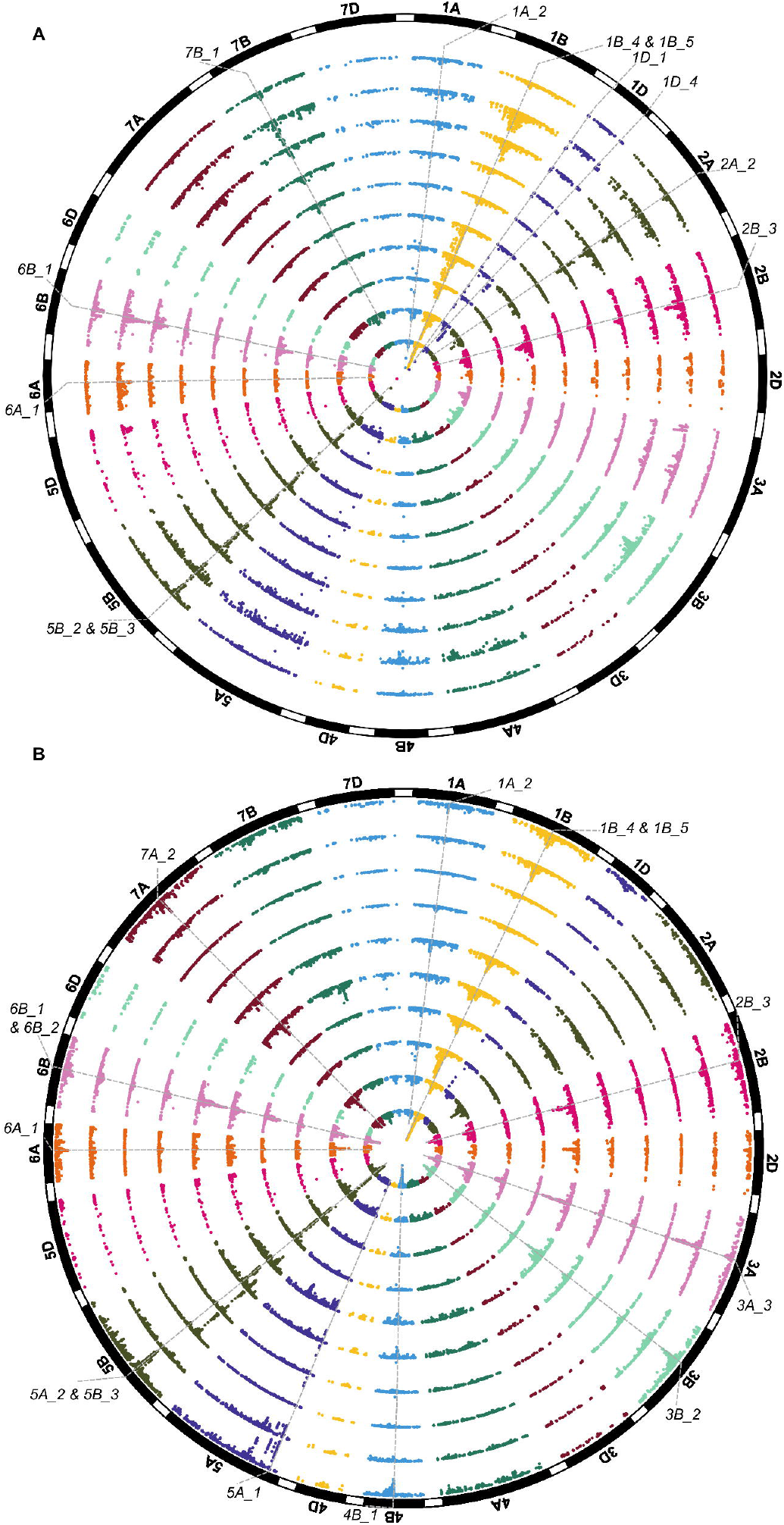
Wheat EigenGWAS for the first ten principal components (PCs). Circular plots of the two wheat panels investigated are shown. Highly significant PCs are in the inner circle and the least significant outer circle are displayed. Genetic positions based on a consensus map (41) are displayed for (*A*) UK winter and (*B*) Brazilian spring wheat panels. Chromosomal introgressions significant across multiple PCs are highlighted (See ***SI Appendix*, Table S9**).

In contrast to wheat, EigenGWAS in the winter and spring barley varieties did not detect any major loci with highly significant peaks across multiple PCs (**Fig. 3** and ***SI Appendix***. PCs variation in **Table S8 &** results in **Table S11**). Although two genomic regions in winter (1H_3 and 4H_3) and three in spring barleys (2H_3, 3H_1 and 7H_1) were identified in at least three PCs. Nevertheless, peaks were also identified close to the locations of known genes controlling flowering time and height (***SI Appendix*. Table S11**), e.g. the PC5 hit on chromosome 3H ∼632 Mbp (explaining 2.46% of the variation) is near the semi-dwarfing gene *sdw1* in spring barley. Interestingly, one of the most significant hits in the spring barley panel (*3H_1*, identified using PC1 and explaining 6.91% of the variation) was also detected using EnvGWAS on variety age and by GWAS on yield (***SI Appendix*. Table S14**). Given the location of this hit in a highly recombinogenic region of the barley genome, and that it was detected only in the spring barley panel, this may indicate a major locus under selection specific to spring barley breeding. No strong peak in winter barley was found for PC1, with the most significant peak obtained using PC6. As UK elite winter barley is more genetically diverse than UK elite spring barley, these results indicate that UK elite winter barley may be subjected to weaker selection pressures. Interestingly, hits on genomic regions (*5H_2* and *7H_1*) from the spring barley EigenGWAS analysis were also identified in GWAS analysis of seasonal growth-habit and variety age, highlighting the importance of these loci under selection (***SI Appendix*. Table 14**).

**Figure 3.**
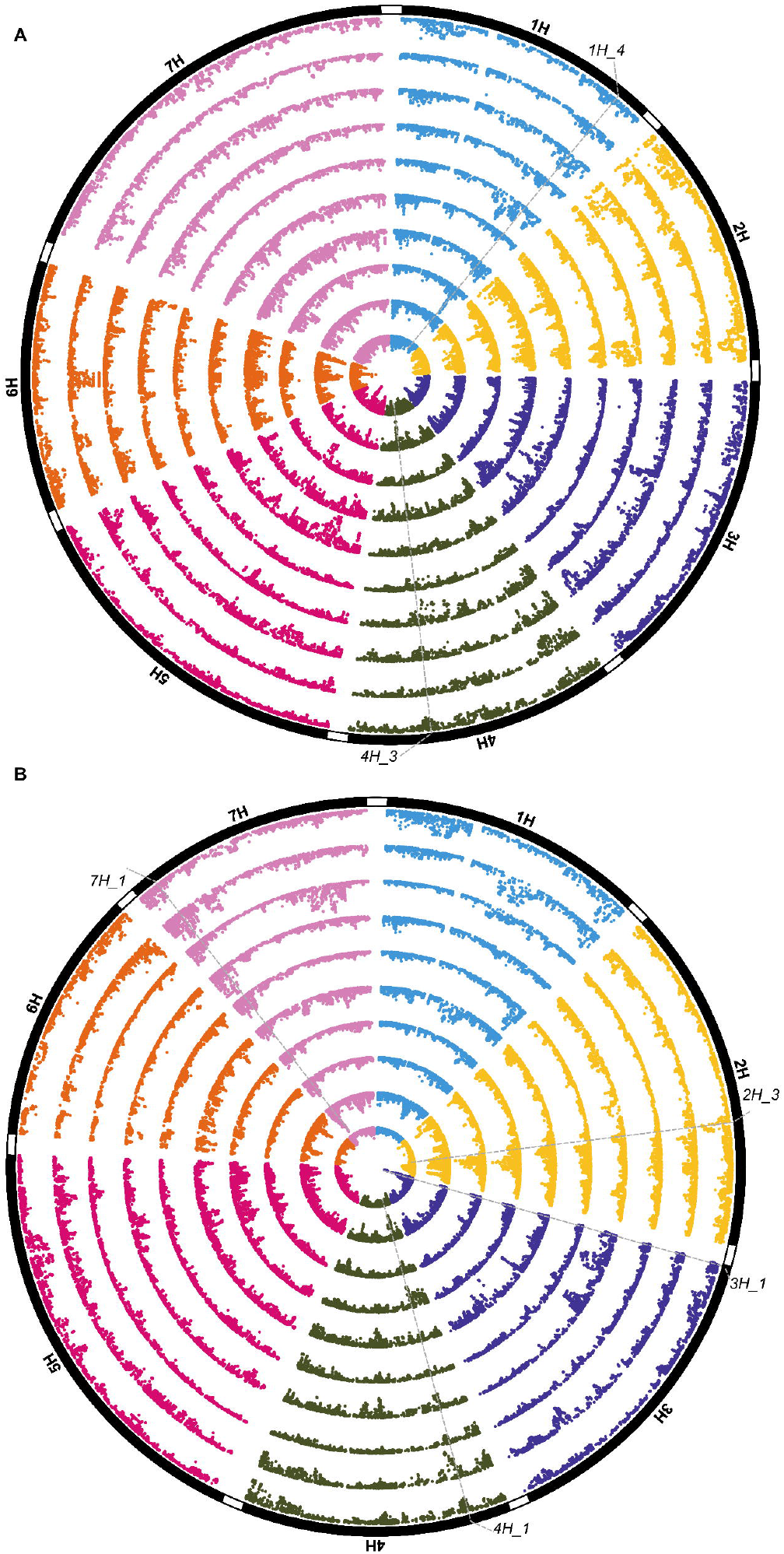
Barley EigenGWAS for the first ten principal components (PCs). Circular plots of the four panels are shown. Highly significant PCs are in the inner circle and the least significant outer circle are displayed. Pseudo-genetic map positions that relate to the physical positions (40) are displayed for (*A*) UK winter and (*B*) UK spring barley panels. Chromosomal introgressions significant across multiple PCs are highlighted (See ***SI Appendix*, Table S11**).

## Discussion

We demonstrate that use of variety age for EnvGWAS can detect regions of crop genomes under selection during breeding. In addition, we show variety age is a good proxy for yield, with the genetic loci identified for wheat validated in an independent experimental multi-founder population (18). Lastly, we showed that the genetic loci detected by EnvGWAS showed gradual, as well as sharp, shifts in allele-frequency over time, indicating subtle changes at these loci by breeders, which are less discernible to detection using approaches such as partitioning the populations on age and searching for differences based on Fst.

It is perhaps not surprising that selection of loci varies between the UK winter and Brazilian spring wheat, given that the target agricultural environments and growth types are very different. Wheat yields in both Brazil and the UK have improved greatly over the years (13, 22). Our contrasting results in wheat indicate that different sets of genes have been selected over the years, and are likely involved in both yield component and local adaptation traits. Future efforts will shed more lights on the types of genes underpinning these loci, allowing changes in allelic diversity over the years to be investigated.

Our results for UK barley contrast with those for UK wheat. Firstly, more hits were associated with variety age in spring compared to winter barley, and secondly an identical peak on chromosome 5H (at ∼19cM, ∼7.5 Mbp) was identified in both panels (as well as in the combined spring and winter analysis). This is surprising as breeders rarely cross spring and winter barley, and since the breeding targets in the two pools differ (malting and largely animal feed, respectively). To further investigate this region, we tested the candidate SNPs against phenotypic data available from national trial data (***SI Appendix*, Table S10**), finding it to be associated with several malting quality traits, powdery mildew resistance and yield in fungicide untreated trials. These findings suggest the potential importance of this region for breeding for disease resistance and end use quality. Interestingly this region on 5H houses a cluster of terpene synthases that have been implicated in fungal disease resistance in other species (23) and that potentially have been selected alongside direct targets such as *Mla* and *mlo* genes (24).

The detection of significant hits with EnvGWAS provides an opportunity to explore their relationship with yield and other agronomically important traits. Some hits coincide with previously published QTL in wheat and barley, for example the highly significant loci on wheat chromosomes 1A and 6A (25–27). Our EnvGWAS hits on chromosomes 1A and 2A also overlapped with the reduced diversity peaks identified in the recent analysis of the UK wheat pedigree by (28). Specifically, the 2A locus may correspond to a stripe rust resistance gene described by (29), as the peak markers overlap. Interestingly a group of R genes *Lr37*-*Yr17*-*Sr38* (30) which were important sources of resistance in the past also lie in this region and might be more plausible candidates, rising in frequency before their resistance broke down. Similarly the highly significant genetic locus on the short arm of barley chromosome 3H for variety age and yield found in the subset of 197 barley lines corresponds to the genomic region associated with the a malting quality trait, hot water extract, in UK spring barley that demonstrated a major change in allele frequency over the last thirty years (31). In addition, the region identified on chromosome 3H (∼68cM) for variety age in winter barley in the larger dataset has been shown previously to be associated with yield component traits (grain length and grain area) in European winter barley (32).

Similarly, in barley, the region identified on chromosome 2H (∼65cM, ∼621 Mbp) for variety life-span has been shown previously to be associated with yield and yield component traits (32, 33) and may correspond to the *OsBR1*/*D61* candidate genes reported previously that are associated with yield traits in barley (32, 33).

This is interesting as old varieties, despite being less-productive than modern varieties, were under cultivation for longer periods. It may however be noted that with the introduction of modern breeding practices yield increases, but with drastic effects on variety life-span due to the more frequent introduction of new varieties that outperform contemporary varieties. In wheat, EnvGWAS on variety life-span also identified a hit on chromosome 1A that co-located with a hit for variety age. This further indicates a direct relationship between variety age and variety life-span in wheat and barley.

Using EigenGWAS, we detected major introgressions in the wheat varietal panels investigated, with several of these found to be in common between the UK winter and Brazilian spring wheat panels, indicating their wide use in breeding. (18), analysing the 16 founder MAGIC population we used in our validation studies here, proposed a major role for multiple introgressions from wild species in UK wheat breeding to date. In contrast, EigenGWAS results in barley provide no evidence of a similar pattern of introgressions in either the winter or spring panels. Wheat and barley breeding differ in their exploitation of genetic resources. In wheat, several alien-introgressions from related species are known to have occurred (34). While wheat is an allohexaploid and can support large tracts of non-recombining alien chromosome, this may not be the case in diploid barley. However, examples of introgressions in barley from landraces and spontaneous mutant lines for agronomically important genes have been reported, such as the semi-dwarfing allele *sdw1d* from the variety Diamant and the disease resistance gene *mlo11* from Ethiopian landraces (24, 35).

Interestingly, within the genomic region of 6A_1, detected by EigenGWAS in wheat (a non-recombining peri-centromeric region) lies the gene *TaGW2* (36) which influences grain-weight and protein content traits that further suggest that the present approach is very effective in discovering genomic regions undergoing selection for yield. Another interesting finding is that the semi-dwarfing *Rht2* gene in wheat (chromosome 2D) was not detected despite its importance in the breeding history of the crop. This could be due to population structure control of the analyses. In the case of *Rht2*, it is noteworthy that GWAS on a panel of French, German and UK lines failed to detect an effect on yield or height unless a locus specific marker was used (37, 38), suggesting weak LD and low marker coverage on the 4D chromosome as the cause of failure here too.

## Conclusion

Breeding has resulted in considerable and sustained genetic improvement of wheat and barley in recent decades, and our results identify at least some of the major loci that have contributed, and are still contributing, to these improvements. Using EnvGWAS, we demonstrate the utility of analysing variety age as a surrogate for traits selected by breeders to detect the genetic loci under selection over time, and to assess the temporal changes in their respective allele frequencies. For UK cereals, trends over time suggest that these loci are likely QTL for yield or yield components. While the resolution of this study in the non-recombining peri-centromeric region is insufficient to definitively associate known QTLs with the loci we have found, several such QTLs were found. EigenGWAS on the same data proved a simple method of detecting contrasting features of genome organisation in wheat and barley, and in some cases these too could be related to traits. We advocate the use of variety age as a surrogate trait, and the use of EnvGWAS and EigenGWAS to identify the genetic loci under selection that have underpinned the productivity gains made via breeding. These extensions to GWAS that exploit historical datasets are useful additions to the analysis toolbox of crop quantitative genetics.

## Materials and methods

### Germplasm, age and trait data

For both wheat and barley, we selected two panels of varieties representing national list entries and some older varieties from the UK (404 winter wheat; 297 winter and 406 spring barleys) and Brazil (355 spring wheat) (**Table S12**). The Brazilian spring wheat panel included entries released between 1922 to 2013. Year of varietal release and trait data were obtained from (39). The UK wheat panel consists of winter wheat varieties that were either registered or in use from 1916 to 2010. The winter and spring barley panels consisted of varieties grown in the UK from 1960 to 2016. Only two-rowed spike morphology types were included and all hybrid varieties were excluded. Variety age for UK germplasm was determined from the year of entry into national list trials or from the first reported year of trial data and was manually checked across different local data and published sources ((13); https://ahdb.org.uk/rl & https://www.gov.uk/government/publications/plant-varieties-and-seeds-gazette-2020 https://www.niab.com/services/seed-certification/botanical-descriptions-varieties) with unresolvable ambiguities removed, reducing the UK wheat panel from 450 to 404 varieties. Following (13), only varieties with either three years trials data or equivalently which were known to be successful in national list trials were included in the dataset. In addition to variety age, we computed life-span of UK varieties as the difference between the last and first year in national trials plus one. This is usually equally to the total number of years each variety remained in trial, though with some rare breaks in the testing sequence over years. Grain yield data for the UK wheat and barley panels were sourced from (13).

### Genotyping

Genotypic data were sourced from NIAB (https://www.niab.com/research/agricultural-crop-research/resources) and JHI (http://www.barleyhub.org/projects/impromalt/) by permission through WAGTAIL and IMPROMALT projects.

For wheat, 14654 SNPs derived from genotyping with the 90K Illumina iSelect SNP array (Wang et al. 2014) generated within the Biotechnology and Biological Sciences Research Council grant BB/J002542/1 were sourced with permission from NIAB, and available at https://www.niab.com/research/agricultural-crop-research/resources. For barley, 43799 SNPs genotyped using the 50K Illumina iSelect array (40) were sourced from (31). Genetic maps for wheat (41) and barley (31, 40) were previously described. The physical map locations of wheat and barley SNPs were retrieved from (42) and (40), respectively. SNPs with a minor allele frequency <5%, missing values <10% and heterozygosity >10% were removed, leaving 12656 wheat SNPs and 25562 barley SNPs for downstream analyses.

### EnvGWAS and EigenGWAS analysis

EnvGWAS and EigenGWAS analyses were performed using the R-package GWASpoly (43) implemented in R version 3.5.2 (http://www.R-project.org/). To determine the population structure of the panels, principal component analysis (PCA) was performed using the R-package SNPRelate (44). The first ten principal components associated with the largest eigenvalues were used for EigenGWAS. Population structure and kinship effects were controlled by inclusion of a mixed model of a canonical relationship (kinship) matrix (45), generated from a subset of SNPs (741 for wheat and 2500 for barley) pruned based on genetic positions. For ease of comparison across GWAS scans, the threshold for significance was set to –log_10_(*p*-value)= 4.0 which in several GWAS scans was above the threshold obtained using false discovery rate (http://www.strimmerlab.org/software/fdrtool/index.html). Manhattan plots and circular plots were generated using R-packages qqman (46) and CMplot (47), respectively.

### MAGIC wheat analysis

The significant SNPs from the wheat EnvGWAS were used for validation against agronomic traits identified in the ‘NIAB Diverse MAGIC’ population (18). Analysis was performed in R using adjustments for the funnel structure of the cross (48). All data used were obtained from the following websites that hosts the genotyping and phenotyping data of the 550 MAGIC-diverse RILs http://mtweb.cs.ucl.ac.uk/mus/www/MAGICdiverse/index.html.

## Data availability

Genotypic data sets of the study are available from NIAB, UCL and JHI via following websites: https://www.niab.com/research/agricultural-crop-research/resources http://mtweb.cs.ucl.ac.uk/mus/www/MAGICdiverse/index.html http://www.barleyhub.org/projects/impromalt/.

## Acknowledgments

This project was part funded by projects IMPROMALT BB/K/0070251/1 and WAGTAIL BB/J002542/1, jointly funded by the BBSRC and UK wheat and barley breeders; and by direct funding of Rajiv Sharma from SRUC. We thank Mark Looseley and Hazel Bull (JHI), for sharing the genotypic data set of the barley varieties analysed here, as well as Chin Jian Yang, Ian Dawson and David Marshall (SRUC) for helpful discussion throughout the work.

## Supplementary Figures

**Fig S1**. Linear trend in variety year of release and yield (t/ha). (*A*) UK wheat; (*B*) UK barley.

**Fig S2**. UK barley Manhattan plots for EnvGWAS from combined winter and spring varieties (*n*=704). *(A)* Manhattan for the age from the analysed barley varieties; *(B)* analysis using seasonal growth-habit (winter and spring-type) as a covariate; *(C)*; GWAS analysis of seasonal growth-habit; *(D)* GWAS analysis of seasonal growth habit with the four peaks identified in *C* as covariates.

**Fig S3**. UK winter wheat Manhattan plots for EnvGWAS validation using a subset of 192 varieties. Manhattan plots are shown for: (*A*) the first year of variety in National trial; (*B*) the last year each variety is enlisted on the national list; (*C*) variety life-span, i.e. how long a variety is on the national list; (*D*) GWAS on yield.

**Fig S4**. UK barley Manhattan plots for EnvGWAS validation using a subset of 197 varieties. Manhattan plots are shown for: (*A*) the first year of variety in National trial; (*B*) the last year each variety is enlisted in the national list; (*C*) variety life-span, i.e. how long a variety is on the national list; (*D*) GWAS on yield.

**Fig S5**. Rocket-plots displaying temporal changes in allele frequencies in wheat (UK-wheat: *A-D* & Brazilian-wheat: *E-H*) and barley panels (winter: *I-J* & spring: *K-N*). Also displayed wheat categories in parenthesis within the panel.

## Supplementary Tables

## Notes

### Competing Interest Statement

The authors have declared no competing interest.

