## Supplementary figures and images for "Trends of genetic changes uncovered by Env- and Eigen-GWAS in wheat and barley"

### Fig. S2.png

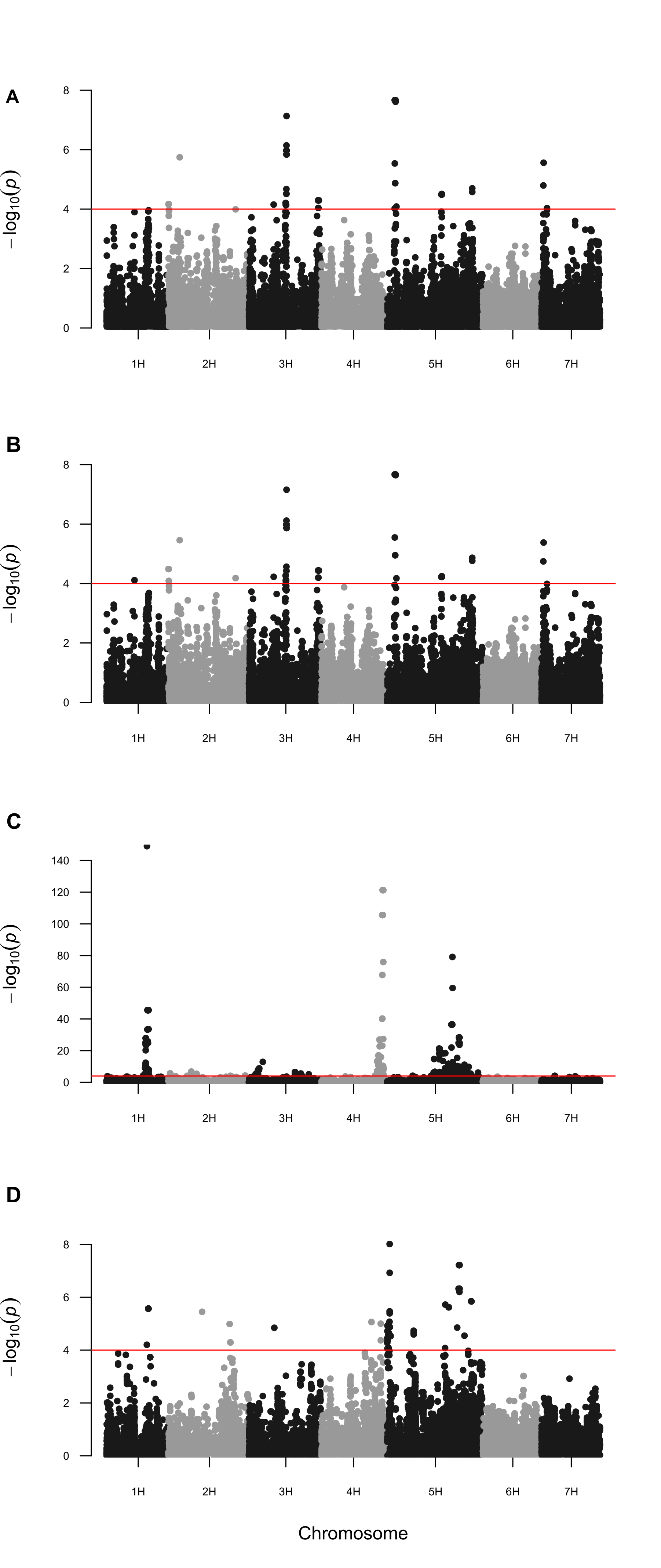

### Fig. S3.pdf

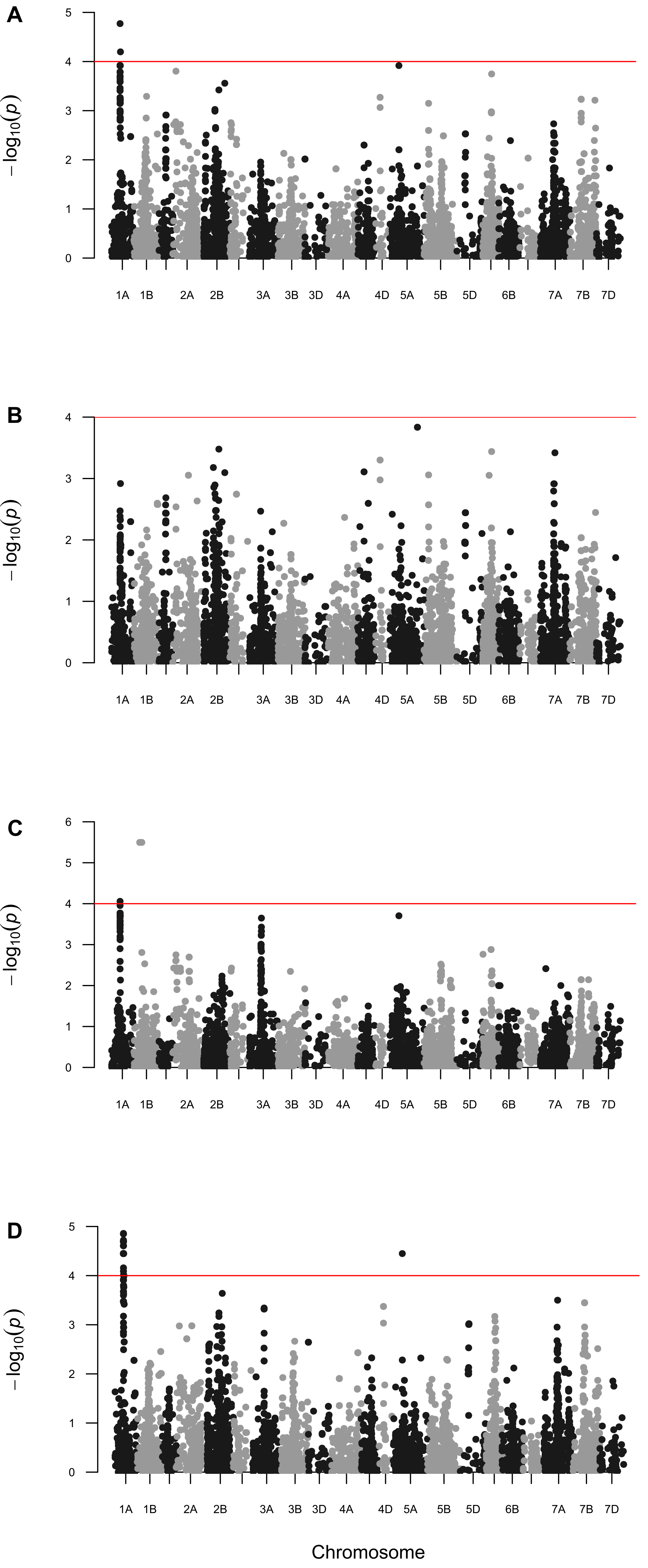

### Fig. S4.png

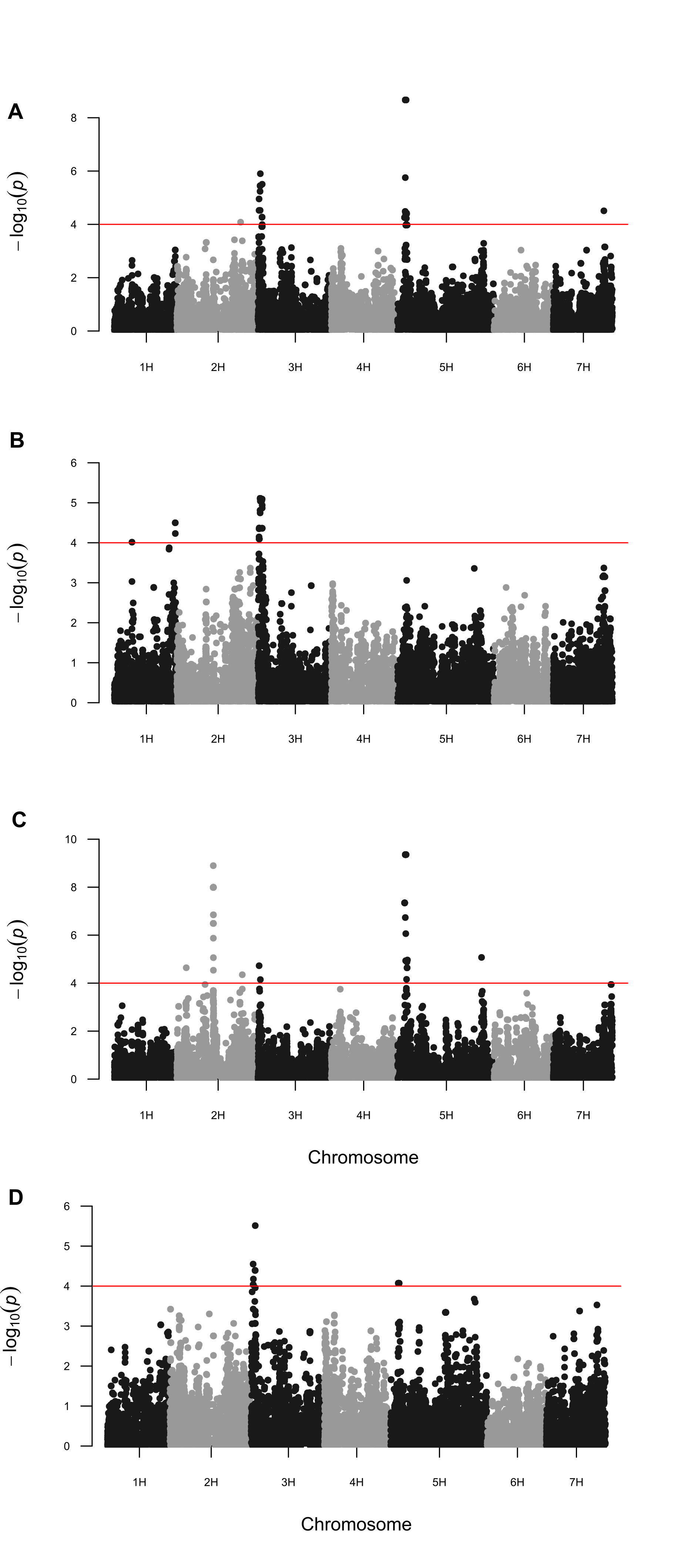

### Fig. S5.pdf

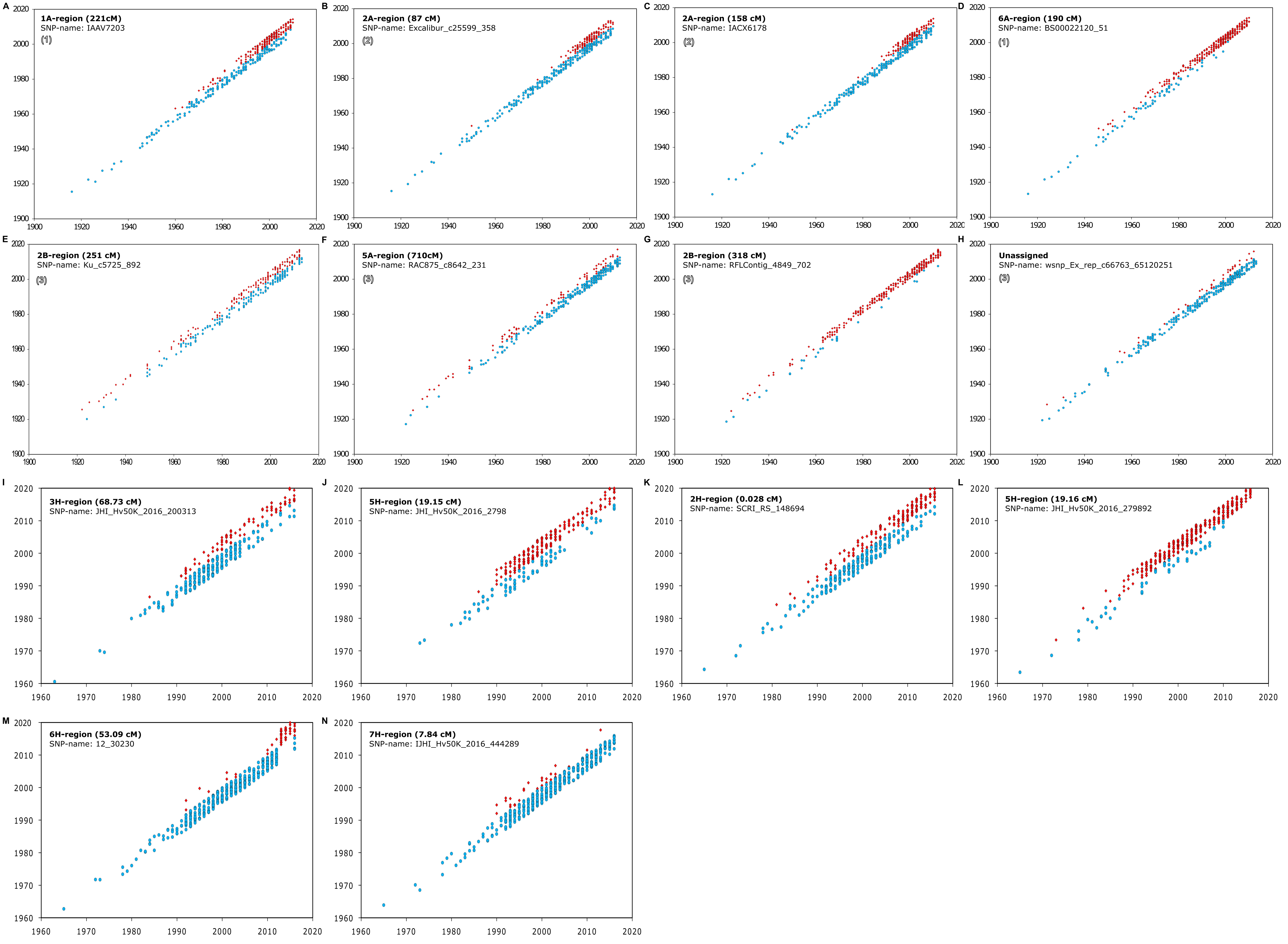
